# Heteroplasmy and repeat expansion in the plant-like mitochondrial genome of a bivalve mollusc

**DOI:** 10.1101/2020.09.23.310516

**Authors:** Andrew Calcino, Christian Baranyi, Andreas Wanninger

## Abstract

**Background:** Animal mitochondrial genomes are typically circular, 14-20 kb in length, maternally inherited, contain 13 coding genes, two ribosomal genes and are homoplasmic. In contrast, plant mitogenomes display frequent gene rearrangements, often contain greatly expanded repetitive regions, encode various open reading frames of unknown function and may be heteroplasmic due to differential repeat expansions between molecules. Error correction by recombination is common in plant mitochondria and has been proposed as the driver behind the rearrangements and repeat expansions that are often observed. In contrast, most animal mitochondria never or only very seldomly recombine and their utilisation of other repair mechanisms for mitochondrial genome error correction is a potential driver of their non-coding DNA reduction.

**Results:** Using PacBio long reads for genome assembly and structural variant detection, we identify evidence of heteroplasmy in the form of variable repeat lengths within two blocks of repetitive DNA within the expanded 46 kb mitochondrial genome of the bivalve mollusc, quagga mussel, *Dreissena rostriformis*. The quagga mussel also has a greatly expanded repertoire of coding genes in comparison to most animals which includes an additional nine open reading frames (ORFs) encoding predicted transmembrane peptides of unknown orthology.

**Conclusions:** The genome size, repeat content and coding gene repertoire of the quagga mussel mitogenome closely resemble those of plants and the identification of repeat-associated heteroplasmy is consistent with the utilisation of plant-like recombination-based error correction mechanisms. Given the frequency of mitochondrial repeat expansions within the Bivalvia, recombination may be an underappreciated mechanism for mitogenomic error correction within this and other animal lineages.

**Significance Statement:** Unlike most animals, the mitochondrial genomes of many bivalve molluscs are often greatly expanded and contain large non-coding regions and additional predicted genes of unknown function. While these features are uncommon in other animal groups, they are common features of plant mitochondrial genomes. Here we show that the mitochondrial genome of the bivalve mollusc, the quagga mussel, displays many plant-like features and additionally, shows evidence of variability in the repeat lengths between mitochondrial molecules within an individual mussel. We propose that similar error correction mechanisms in plants and bivalves may play a role in these observed commonalities.

## Introduction

The traditional view of animal mitochondrial biology is one of streamlining of non-coding DNA, strict maternal inheritance and homoplasmy. For mammals and many other model species, this generalisation holds true with both paternal inheritance and heteroplasmy associated with pathological conditions. One of the most dramatic deviations from this scenario comes from a growing list of bivalve mollusc species in which inheritance of unique maternal and paternal mitochondrial isoforms are associated with sex determination and gonad differentiation (Breton et al. 2007; Passamonti and Ghiselli 2009; Zouros 2013). The male and female mitogenomes involved in this system of Doubly Uniparental Inheritance (DUI) can differ by as much as 50% at the nucleotide level and each are defined by the presence of unique male or female specific open reading frames (Zouros et al. 1994; Breton et al. 2009).

Outside of the traditional model species, the flood of new mitochondrial genomes being sequenced with the aid of modern high throughput technologies has revealed mitochondrial biology to be a much more diverse and complex landscape than commonly appreciated (Breton et al. 2014). The longest mitochondrial genome yet sequenced belongs to the cnidarian tube anemone *Isarachnanthus nocturnus* at 80,923 bp, which is linear and divided over five chromosomes (Stampar et al. 2019). Within the molluscs and particularly within the bivalves, genome expansions and rearrangements are also commonplace. The largest molluscan mitochondrial genome sequenced to date belongs to the zebra mussel *Dreissena polymorpha* at 67 kb (McCartney et al. 2019) while a recent study has highlighted frequent expansions in the ark shells with the longest reported mitogenome coming from *Scapharca kagoshimensis* at 56 kb (Kong et al. 2020).

The size of these expanded genomes are remarkable amongst the animals, however they are frequently dwarfed by those of plants. The Sitka spruce, for example, has a mitogenome size of 5.4 Mb and encodes 41 protein-coding genes (Jackman et al. 2020). Although DUI is restricted to a subset of bivalve lineages, mitochondria-based sex determination mechanisms have also evolved in plants through gynodioecy, in which females are derived from the hermaphroditic condition through cytoplasmic male sterility (CMS, Chase 2007). In all cases known, CMS is brought about by mutations in mitochondrial genes (Hanson and Bentolila 2004). While exploring parallels between the DNA mismatch repair protein MutS in soft corals and plants, (Abdelnoor et al. 2006) noted several convergent properties of the mitochondrial biology of the two clades and suggested a portentially expanded role for mitochondrial activity in sex determination of both that might be related to their sessile lifestyle. The vast majority of bivalve species have a sessile or semi-sessile lifestyle as adults and it is striking that within this group, the sex-linked DUI condition evolved.

Here we investigate the genomics and transcriptomic dynamics of the mitochondria of the quagga mussel, *Dreissena rostriformis*. Freshwater dreissenid mussels evolved from marine ancestors during the late Miocene period (Harzhauser and Mandic 2010). Unlike the freshwater unionid mussels which have evolved a derived parasitic larval stage known as a glochidium (Watters 1999), dreissenids have retained the indirect mode of development common to many marine species which includes lecithotrophic trochophore and planktotrophic veliger stages (Cragg 1996; Giribet 2008). Dreissenids belong to the Imparidentia (González et al. 2015; Combosch et al. 2017) and are important invasive colonisers of freshwater habitats in Europe and North America (Mills 1993; Heiler et al. 2013).

Although the nuclear genome of the quagga mussel was recently published, the mitochondrial genome was not resolved, most likely due to the large repetitive regions which could not be spanned by Illumina short reads (Calcino et al. 2019). We reveal the presence of heteroplasmy resulting from variable repeat expansions within an individual and not due to DUI as is common in many bivalve species. We discuss the parallels between plant and bivalve mitochondrial genome organisation and suggest that future studies should focus on DNA repair processes within the Bivalvia, in particular with the utilisation of homologous recombination for double strand break repair.

## Results

### Genome assembly and gene annotation

The mitochondrial genome assembly of the quagga mussel spans 46,868 nt and encodes all 13 of the coding genes typical for metazoan mitochondria (Fig.1). The large and small mitochondrial ribosomal subunits are also present, as are tRNA genes for all 20 amino acids including two leucine, three arginine, two serine and two tyrosine copies (Supplementary Table S1). Additionally, a predicted suppressor tRNA with an anticodon corresponding to the stop codon UAA was identified. All protein-coding genes start with either an ATG codon (six genes) or an ATA codon (seven genes).

**Figure 1.**
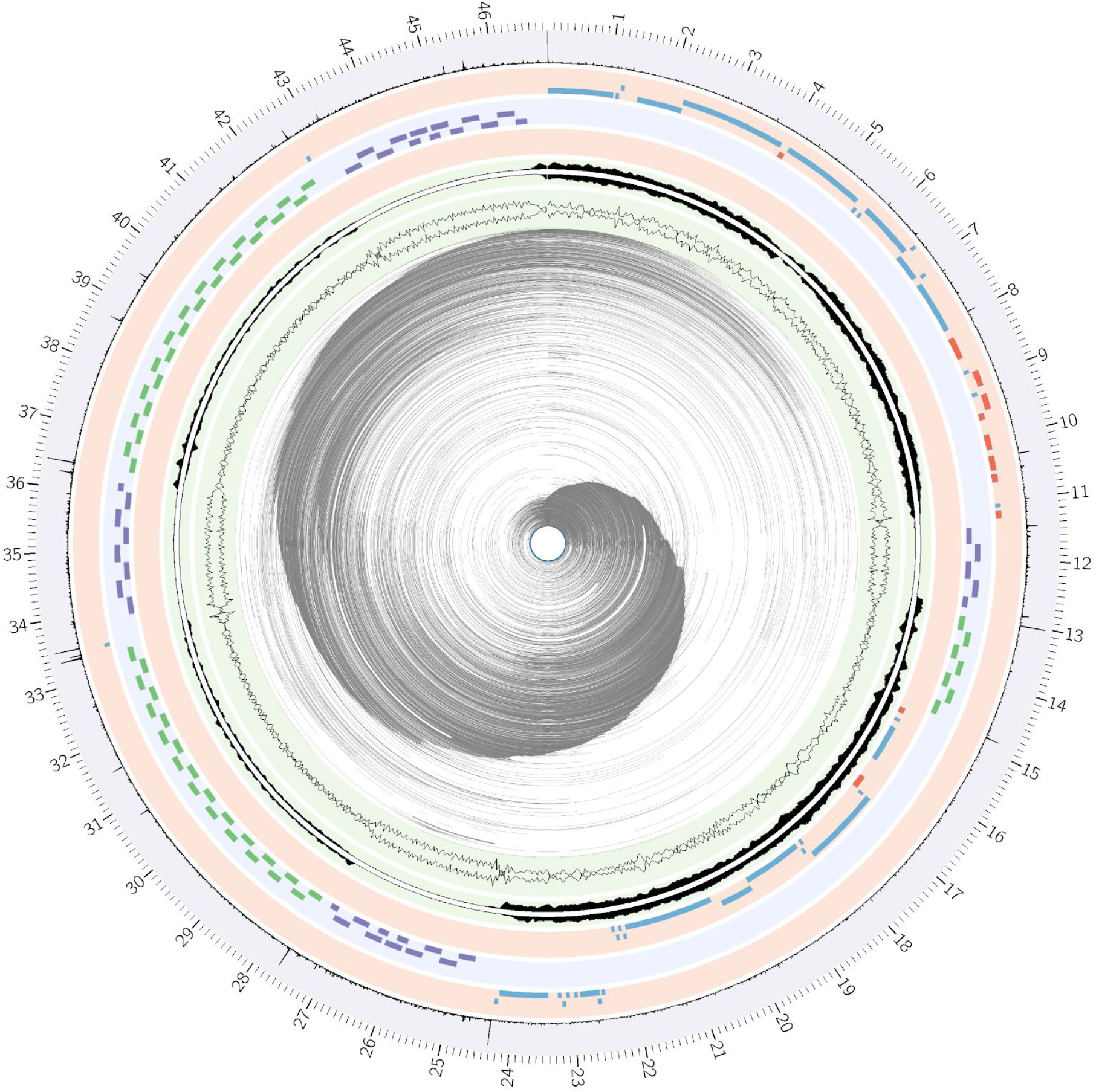
The mitochondrial genome of the quagga mussel. Starting from the innermost track, grey lines represent mapped PacBio reads generated by ConcatMap. Black wiggle plot shows GC/AT content using a 40 bp sliding window. Black histogram shows short read coverage on the heavy strand (facing out) and the light strand (facing in). Coding and ribosomal genes (long, blue), tRNAs (short, blue) and novel ORFs (red) map to the light strand. 308 bp (purple) and 258 bp (green) tandem repeat loci. Coding and ribosomal genes (long, blue), tRNAs (short, blue) and novel ORFs (red) map to the heavy strand. Histogram of mapped PacBio read start and end sites.

Compared to the congeneric zebra mussel *Dreissena polymorpha* (Soroka et al. 2018; McCartney et al. 2019), the quagga mussel mitochondrial genome appears to have undergone a single major rearrangement in which the region encoding nad4l and cox3 has been transferred from its position between cox2 and atp6 to the region downstream of nad5 on coding block 2 (Fig. 2).

**Figure 2.**
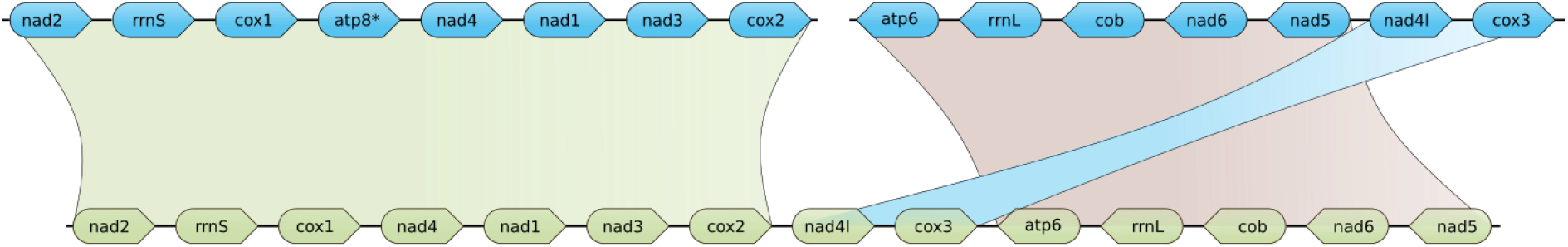
Comparison of gene arrangement in the quagga mussel (above) with the zebra mussel (below). A single translocation of the quagga mussel nad4l and cox3 relative to the zebra mussel is depicted. atp8* was not identified in the zebra mussel (Soroka et al. 2018; McCartney et al. 2019). The gap between cox2 and atp6 in the quagga mussel represents the repetitive region between coding blocks 1 and 2.

Similarity between the quagga mussel gene and amino acid sequences with those of the zebra mussel reveal a high degree of similarity with the only exception being the 5’ region of nad1 which is shorter in the quagga mussel. Amino acid sequence alignment of the region between the putative nad1 start sites of the two species (NCBI: APX39121.1) shows high similarity except for an alanine/valine at position seven and two in-frame methionines in the zebra mussel at positions 1 and 8 which correspond to an alanine and a leucine, respectively, in the quagga mussel (Supplementary Figure S1). No in-frame methionine is encoded 5’ of the putative start site in the quagga mussel before reaching a stop codon located at nucleotide position 5673-5675. To confirm that this was not the result of a sequencing or assembly error, short Illumina reads corresponding to the 5’ start site of nad1 were extracted from datasets from a second individual (SRA BioSample SAMN12125624, Supplementary File S1). The result was a perfect alignment between the predicted amino acid sequences from these samples with the one presented here.

Atp8 was not detected by Mitos2. However, one of the 11 ORFs detected, ORF1, which is located on the heavy strand between cox1 and nad4 at nucleotide position 3998-4091, bears resemblance with other Imparidentia atp8 genes (Supplementary Figure S2). Atp8 and the remaining 10 ORFs all possess between one and four predicted transmembrane domains. However, BLASTp and SWISSMODEL revealed no insightful or significant similarity for any of the sequences (Supplementary Table S2).

### Repeats, gc content, coverage and clipped read ends

The coding genes including all ten uncharacterised ORFs and 24 of the 26 tRNAs lie within two blocks that are separated by large stretches of non-coding DNA. This non-coding DNA is composed of seven blocks of two varieties of tandem repeat which are 258 nt (three blocks) and 308 nt (four blocks) long (Fig. 1). These tandem repeat elements will be referred to as tr258 and tr308 henceforth. The consensus sequence of tr258 and tr308 do not align and although the arrangement of large tandem repeat expansion has also been observed in the zebra mussel (McCartney et al. 2019), the length of the repeats (125 bp, 1030 bp, and 86 bp in the zebra mussel) is dissimilar to those in the quagga mussel.

GC content of the mitochondria as a whole is 33.1% but this value varies significantly between the coding regions and the two repeat elements. In coding block 1 (position 1-11302) gc content is 33.5%, in coding block 2 (position 14948-24302) it is 31.3%, across all copies of tr258 it is 23.6% and across all copies of tr308 it is 46.7% (Fig. 1).

Mapped short read coverage also varies between the two coding blocks and the two repeat elements (Fig. 1). This occurred due to the filtering of the data prior to mapping against the mitochondrial genome by first removing reads that mapped to the nuclear genome. The highest coverage was observed for the two coding blocks followed by tr258 and tr308. To determine if these repeat elements also exist in the nuclear genome, the unfiltered libraries were first mapped against the mitochondrial genome with reads mapping to the repeats extracted. These were then mapped against the nuclear genome. Most of the reads that mapped to both the nuclear genome and the repeat elements of the mitochondrial genome had very short alignments with nuclear sequences. The majority of nuclear alignments were under 30 nt in length and corresponded to low complexity regions of the repeat elements typically with a high nucleotide bias. No evidence for the existence of tr258 or tr258 in the nuclear genome was uncovered.

The ConcatMap figure of PacBio reads mapped to the genome assembly showed evidence that particular sites within the assembly appeared to have a higher than expected density of mapped read start and end sites (Fig. 3a). When clipped ends are retained it becomes clear that many of the mapped reads have large unmapped portions at one or both ends which may be interpreted as evidence of assembly error (Fig. 3b). To ascertain whether or not this was the case, another plot including only reads >40kb was produced (Fig. 3c). This revealed many reads spanning almost the entire assembly, providing evidence that the assembly indeed did reflect the actual sequence for at least some of the mitochondrial DNA molecules within the sample.

**Figure 3.**
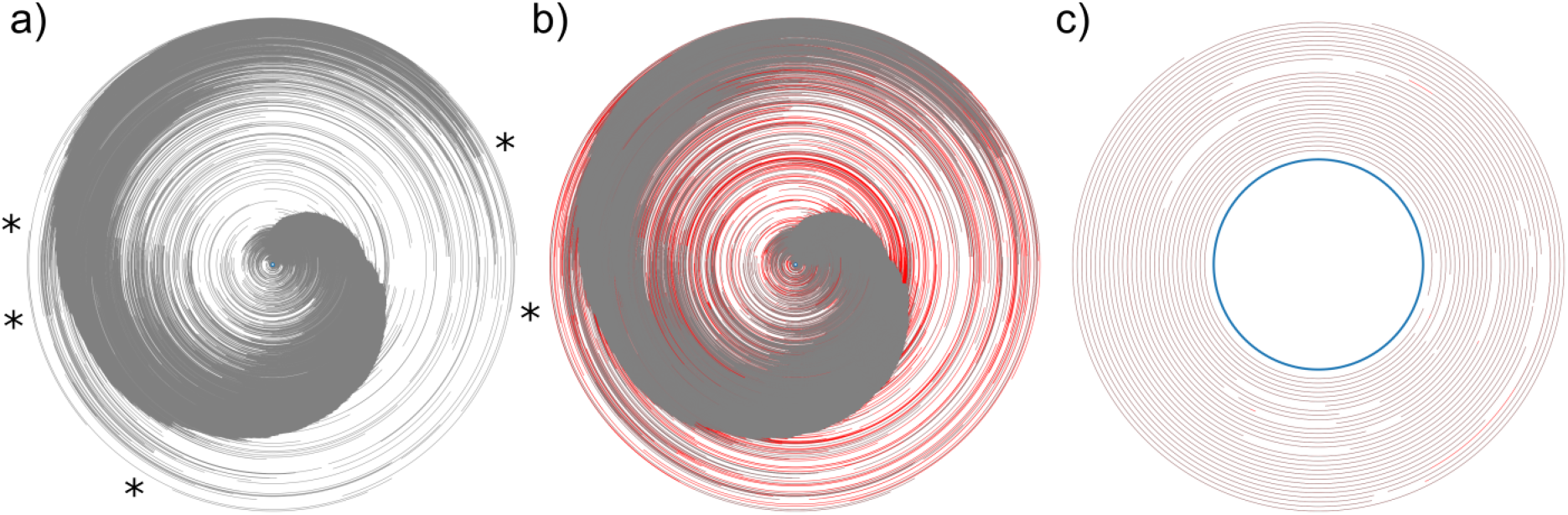
Concat maps of the mitochondrial genome assembly. a) All reads greater than 5kb in length with unmapped ends clipped. Five loci that display a high density of mapped start and end sites are marked with a star (*). b) All reads greater than 5kb in length with unmapped ends retained (red). c) All reads greater than 40 kb in length.

After extracting reads that had been clipped between nucleotide positions 33,000 and 34,000, the unmapped ends of each were themselves aligned to the genome, independent of the original mapped regions of the reads. Most clipped ends mapped to one of the two coding blocks, indicating that perhaps the initial alignment had assigned them to the incorrect repeat block. Despite this, ten clipped ends aligned entirely to the large repeat block and the full length reads from which these clipped ends derived were further investigated for repeat content (Supplementary Figure S3).

In each of the ten reads, the copy numbers of tr258 and tr308 were variable, however the most variable repeat was a previously undetected ∼67 bp (tr67) repeat element (Supplementary File S3). Re-analysis of the tandem repeats from the genomic sequence corresponding to this region showed that, while tr67 was present in the assembly, it overlapped tr308 almost entirely and so was not annotated as an independent repeat element (Fig. 1, Supplementary File S2). The copy number of tr67 corresponding to this region ranged from 2.7 in the genome assembly to at least 22 in one particular read (the reasons for the inexact number are due to the error rate of PacBio reads, partial copies expressed as fractions and the potential overlap with tr308 elements).

### Transcriptional dynamics over developmental time

Gene transcript abundance, as measured by transcripts per kilobase million (TPM), can vary by over four orders of magnitude within a single developmental stage (Fig. 3). This is exemplified by the D-shaped veliger 5 larval stage in which the TPM values of ORF 6 and rrnL are 1.67 and 26,778, respectively. If only considering the 13 standard mitochondrial protein coding genes, the lowest and highest abundance genes are nad3 and cob that have TPM values of 18.3 and 13,424, respectively. Although transcript abundance varies significantly between genes, there is clear consistency in which genes are highly expressed and which genes are lowly expressed across developmental time points. For example, nad3, nad4l and several of the novel ORFs consistently display low transcript abundance in all developmental time points whereas rrnL, cob and nad6 are consistently highly abundant (Fig. 4).

**Figure 4.**
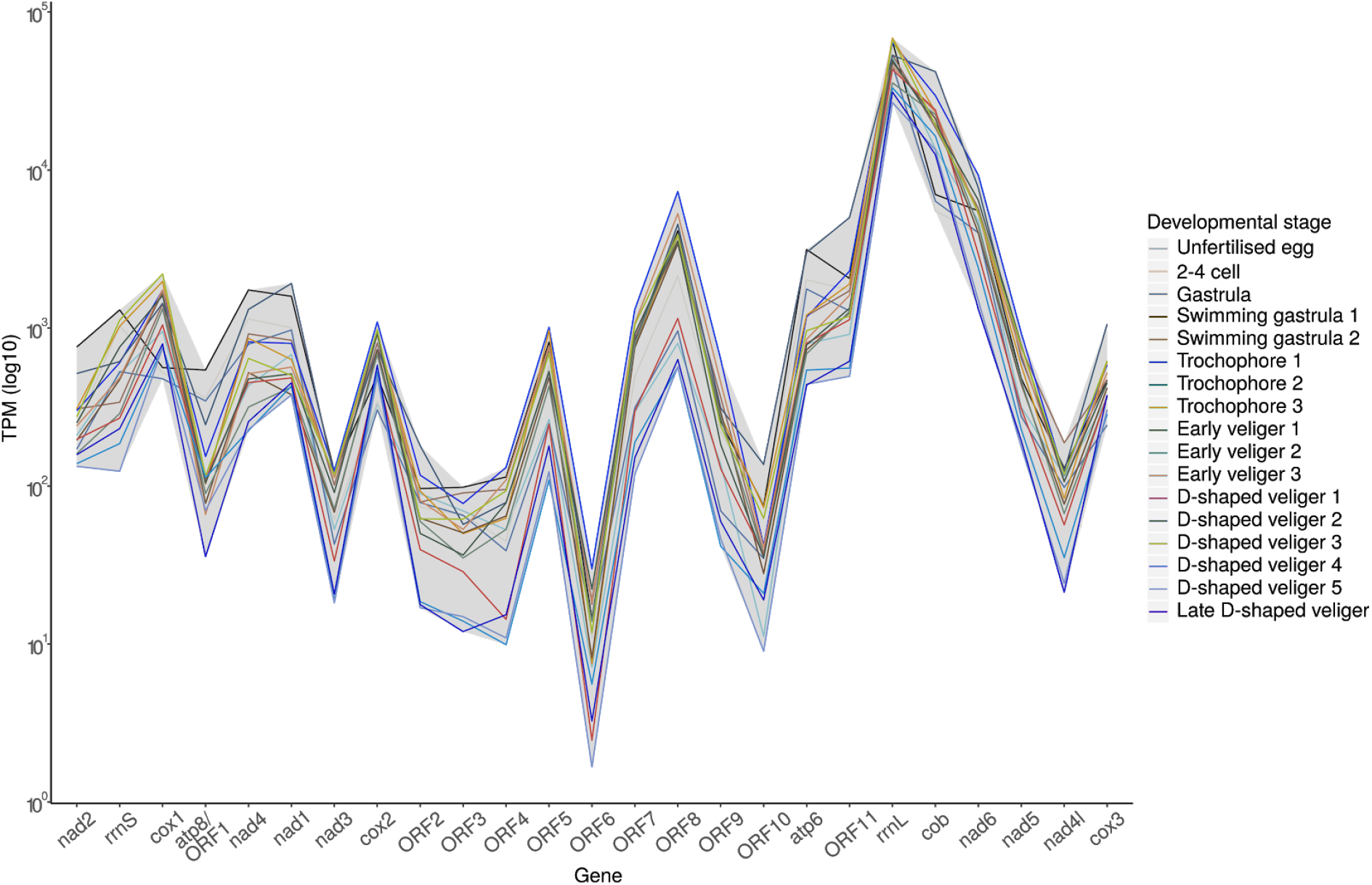
Developmental expression values of mitochondrial genes. Y-axis shows the log of the transcripts per kilobase million (TPM) value and the X-axis displays each of the 13 standard coding genes, the two ribosomal RNAs and the novel ORFs in the order on which they reside on the mitochondrial genome. The grey region highlights the lower and higher range of abundance for each gene.

To determine if mitochondrial gene expression displays patterns related to developmental time point, to the strand on which they are encoded (heavy or light) or to which of the two blocks of genes that they reside within, a heatmap was produced in which the developmental TPM values are normalised for each gene (Fig. 4).

No clear pattern of expression was observed relative to the location or orientation of the genes. However, the relative abundance of gene transcripts did display some consistency over the course of development. Most notably, transcript abundance levels of most genes appear to peak during the early trochophore stages and then decline towards the later veliger stages.

To visualise RNA-seq expression, circos plots with mapped RNA-seq abundance histograms were produced for each developmental stage (Supplementary Figure S4). These demonstrate consistent expression patterns in which coverage is high for the coding strand with some residual coverage of the template strand also present in both coding blocks. Although multimapping of reads was allowed, very little RNA abundance is detected from the repetitive regions of the genome, however peaks of RNA expression are evident at the boundaries of all repeat blocks.

The search for NUMTs revealed two positive partial hits for NAD1 and NAD4 that mapped to scaffold6889 from the nuclear genome. Manual inspection revealed that the two hits corresponded to the terminal regions of a single 1,326 nucleotide DNA block that aligned to the mitochondrial genome from position 4,519 to 5,916 (Supplementary File S3) using MAFFT with settings --maxiterate 1000 --localpair (Katoh and Standley 2013). The longest ORF predicted from the NUMT using the standard genetic code was 79 amino acids long and had a top BLAST hit to a 50 amino acid region of the 463 amino acid long nad4 protein of *D. polymorpha* (APX39122.1).

## Discussion

### The plant-like character of the quagga mussel mitogenome

The mitochondrial genome of the quagga mussel shares several characteristics with plant mitochondria which are uncommon in most animal lineages. The ∼46 kb molecule is one of the largest animal mitogenomes yet described and this genome expansion is the result of two large variable repeat regions which separate the coding genes into two distinct blocks. While near complete coverage of the genome by a number of particularly long PacBio reads provides critical evidence for the accuracy of the genome assembly (Fig. 3c), the discovery that the mitochondrial population of the sequenced individual was heteroplasmic means that it is highly likely that mitochondrial molecules larger than the reported 46 kb assembly were also present in the individual. An obvious question that arises from this is what size range do the mitochondrial genomes of quagga mussels occupy? It is quite likely that sampling of other individuals or the use of different assembly methods would result in the discovery of mitogenomes significantly larger, and perhaps smaller than the reported 46 kb assembly, and that these differences would be the result of variable repeat expansions. Repeat expansions and the existence of multiple mitochondrial genome isoforms within an individual are known from plants but have not been reported for animals (Kozik et al. 2019).

Plant and animal mitochondria differ in size but also in mutation rate. The high coding gene silent site mutation rates of animals relative to plants sits in contrast to the variable, but consistently high, non-coding DNA content of plant mitogenomes (Christensen 2013). To account for the discrepancy of low coding DNA mutation rate and high non-coding DNA mutation rate in plants, Christensen (2013) proposed what has since been termed the “Jekyll and Hyde error correction method” (Smith 2020), in which single stranded DNA damage is converted to double stranded breaks before being corrected by homologous recombination based gene conversion. Jekyll and Hyde refers to the outcomes of this process in which selection against errors in coding regions created by the recombination process would result in only faithfully repaired mitogenomes being retained and therefore observed, while the absence or reduction of selection against errors within repetitive regions would result in slippage and repeat expansion.

### Bivalve mitogenomes

The quagga mussel encodes several ORFs of unknown orthology or function and it has also undergone at least one major rearrangement since the species diverged from the congeneric zebra mussel (Fig. 2). Both novel ORFs and gene rearrangements are commonly found in plant mitogenomes (Christensen 2014) and evidence suggests that these features might also be common amongst bivalve molluscs (Boore et al. 2004; Wu et al. 2012; Breton et al. 2014). A recent study into the mitochondrial architecture in ark shells found that tandem and inverted repeats may be responsible for genome expansions in this bivalve order in which genome sizes range from 17 kb to 56 kb (Liu et al. 2013; Kong et al. 2020).

In species with maternally inherited homoplasmic mitochondria, the detection of recombination is difficult because the invariability of native and recombinant DNA masks such events. An exceptional feature of some bivalve mitochondria is the presence of the Doubly Uniparental Inheritance (DUI) mechanism in which distinct maternal and paternal mitogenomes are simultaneously present within male individuals (Zouros et al. 1994; Zouros 2013; Plazzi and Passamonti 2019). It was in such a non-pathological heteroplasmic system that direct observation of mitochondrial recombination in an animal first occurred (Ladoukakis and Zouros 2001). This observation was possible because the male and female mitochondrial forms of DUI in bivalves can differ by as much as 50% (Doucet-Beaupré et al. 2010), making recombinant mitochondrial molecules clearly distinguishable from both male and female progenitors (Ladoukakis et al. 2011).

Despite the recovery of 10 mitochondrial ORFs of unknown function or orthology, no evidence of DUI was found for the quagga mussel. It also remains to be seen whether or not the silent site mutation rate is typical of other animals. Although the Jekyll and Hyde model predicts a low silent site mutation rate coupled with a high non-coding mutation rate, it is conceivable that a more incremental increase in the role of recombination for mtDNA repair in dreissenid mussels could result in expanded repetitive regions without drastically decreasing the silent site mutation rate. While the size of the mitogenome of the quagga and zebra mussels are exceptionally large for animals, they would be considered on the smaller end of the spectrum for plants.

### Mitochondrial recombination and Doubly Uniparental Inheritance in animals

Animal mitochondrial recombination has now been demonstrated on numerous occasions across multiple phyla, but it remains unclear how commonly these events occur and to what extent recombination is utilised as an error correction mechanism (Alexeyev et al. 2013). Indeed, in mammalian mitochondria, recombination can have serious detrimental effects, particularly in aged tissue, through the production of DNA deletions (Chen 2013). The most well established mitochondrial DNA damage repair mechanisms in animals are DNA mismatch repair (MMR), base excision repair (BER) and non-homologous end joining (NHEJ), which are utilised for repairing small nucleotide mismatches, oxidative DNA lesions and single-stranded DNA breaks as well as double stranded breaks, respectively (see Alexeyev et al., 2013 for a review). The routine utilisation of homologous recombination for double stranded break repair has not been reported in animals, unlike in plants where it appears to be a dominant mechanism for correcting a variety of DNA damage types (Davila et al. 2011; Christensen 2014).

Transcription of the two coding blocks does not appear to be greatly impacted by the interruption caused by the two large repetitive insertions. Although RNA-seq data shows low coverage of the repetitive regions (Supplementary Figure S4), the genes hosted by the two coding blocks show similar patterns of expression that peaks towards the early trochophore stage and dips towards the later veliger stages (Fig. 5). This could be a reflection of increased mitochondrial activity during the transition from the swimming gastrula to the trochophore stage, however the low rate of transcription in the later veligers may also be the result of starvation or other stress due to the lab environment in which they were reared. Regardless, the shared pattern of high and low expression across development by genes on the two coding blocks suggests that the quagga mussel mitochondria are robust to large-scale rearrangements.

**Figure 5.**
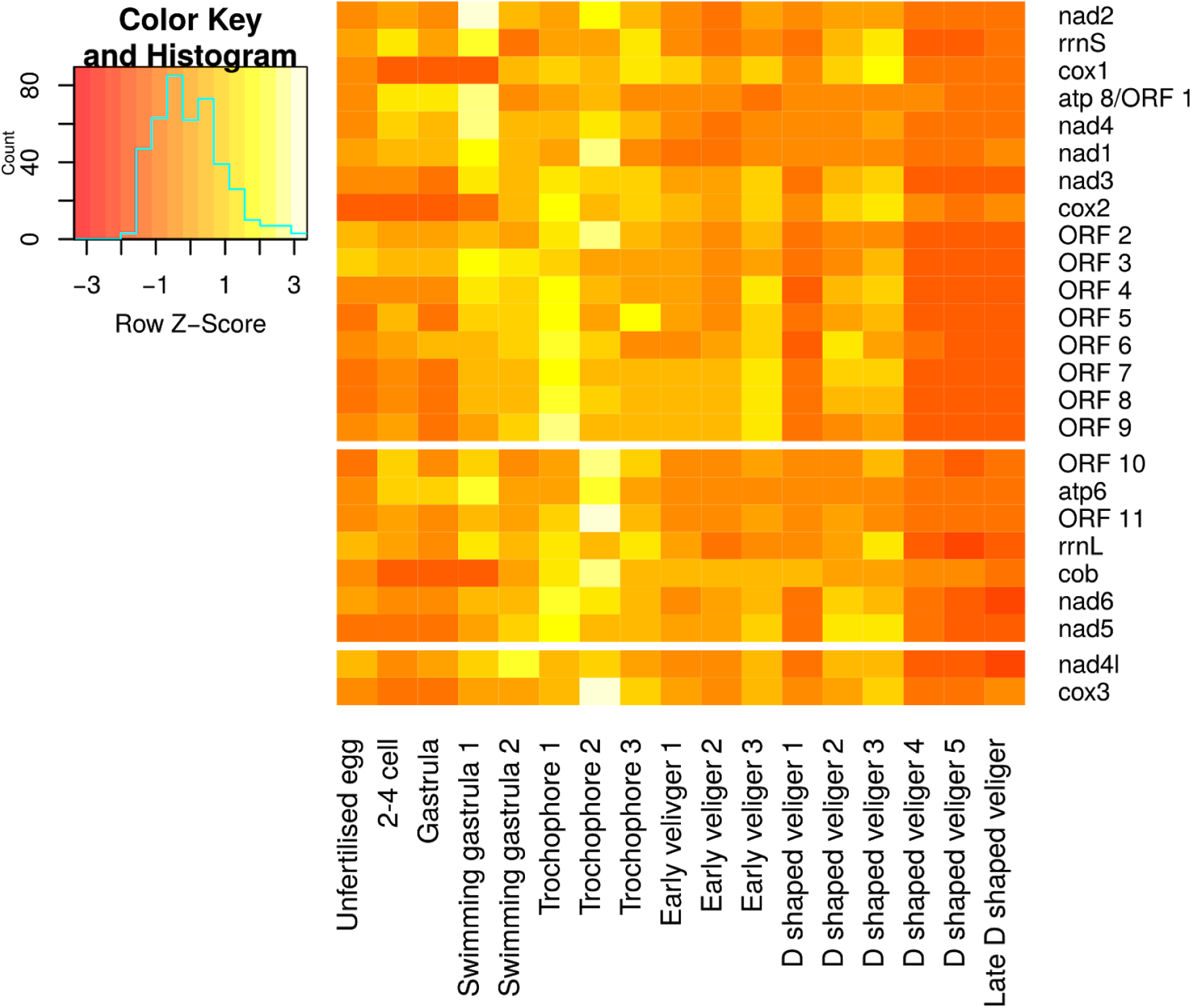
Developmental gene expression heatmap. Each row of TPM values is normalised (Z-score) and the genes are separated into three blocks. The upper block represents all genes from coding block one, all of which are on the heavy strand. The middle block represents genes from coding block two on the light strand and the lower block represents genes from coding block two on the heavy strand.

## Conclusions

We have demonstrated that the mitochondrial population of the quagga mussel *Dreisenna rostriformis* does not consist solely of a single genomic isoform. Rather, two large variable repeat regions are responsible for a mitochondrial population which most likely occupies a range of lengths within any one individual. The plant-like features of the quagga mussel mitogenome add to the list of parallels identified between plant and bivalve mitochondrial biology (Breton et al. 2014) and we hypothesise that some of these observations may be due to similarities in recombination-based error correction methods utilised by the two lineages. With increasing focus on the genomics and energetics of dreissenid mussels, and with continuing advances in sequencing technology, future efforts may aim at documenting the entire range of mitochondrial diversity in this important invasive species.

## Methods

Details of the precise commands used for each of the following steps can be found in the supplementary methods.

## DNA extraction

A single male *Dreissena rostriformis* was collected from the Danube River in Vienna, Austria, and maintained for seven days in 0.2 µm filtered river water. A 300 mg sample of mantle tissue was cut into small pieces on a pre-cooled petri dish, then ground to a fine powder in liquid nitrogen with a pre-cooled mortar and pestle. The powder was then transferred to a 50 ml screw-cap tube and genomic DNA was extracted using the QIAGEN Blood and Cell Culture DNA Maxi Kit (#13362) following the manufacturer’s instructions for tissue samples. The DNA sample was then sent to the Vienna BioCenter Core Facilities (VBCF) sequencing facility for large insert library construction and sequencing on a PacBio Sequel sequencer.

### Genome assembly

An initial genome assembly was produced from canu (v1.8, Koren et al. 2017) corrected PacBio long-read sequences with Genseed (Alves et al. 2016) using the cytochrome oxidase I (COI) sequence as a seed. Long reads were then mapped back to the resulting assembly with pbmm2 (v1.0.0, https://github.com/PacificBiosciences/pbmm2) and an assembly was performed with canu (Koren et al. 2017) using these reads. A second round of pbmm2 mapping was performed on the resultant canu assembly and the assembly polished with gcpp (v1.0.0-1807624).

### Genomic characterisation

GC- and AT-content of the genome assembly was measured using a 40 bp sliding window (https://github.com/DamienFr/GC_content_in_sliding_window). To confirm coverage of the assembly, a short read library (SRR9588035) was first mapped to the nuclear genome with bwa-mem (v0.7.16a-r1181, Li 2013) to filter non-mitochondrial reads, followed by extraction of the remaining paired reads with Fastq-pair (v0.3, Edwards and Edwards 2019) and mapping of these to the mitochondrial genome. Coverage files for production of wiggle plots were produced with bedtools genomecov tool (v2.29.0, Quinlan and Hall 2010). Repeats were identified with Tandem Repeat Finder (v4.09, Benson 1999) following manual inspection to identify the consensus sequence of the largest repeat blocks. These consensus sequences were then used as queries for BLASTn (v2.8.1+, Camacho et al. 2009) searches of the genome.

ConcatMap (v1.2, https://github.com/darylgohl/ConcatMap) was used to visualise long read mappings to the genome. As per the instructions of ConcatMap, a 2x construction of the genome assembly was produced and to this the long reads were mapped with minimap2 (Li 2018). The resulting sam file was used as input to ConcatMap. A second ConcatMap figure was also produced by sampling all mapped reads greater than 40 kb in length. Start and end sites of mapped long reads were extracted with bedtools genomecov tool to identify any indication of heteroplasmy within the sampled mitochondrial population. To determine what differences were present in the clipped reads versus the assembled sequence, full length reads clipped between nucleotide positions 33,000-34,000 and which had their clipped ends re-mapped back to repetitive loci were extracted and investigated for tandem repeat content with Tandem Repeat Finder.

### Gene annotation and transcriptional dynamics

Coding genes and tRNAs were annotated using Mitos2 (Donath et al. 2019) followed by manual annotation to confirm the correct 5’ and 3’ ends of the genes. To annotate non-standard mitochondrial open reading frames (ORFs), developmental de novo transcriptomes were built for the 18 datasets (PRJNA551098) used to annotate the quagga mussel nuclear genome (Calcino et al. 2019) using Trans-ABySS (v2.01, Robertson et al. 2010). All de novo assemblies were combined and deduplicated with cd-hit (Li and Godzik 2006) and these were mapped against a genome assembly with minimap2 that included the nuclear and mitochondrial scaffolds. All transcripts that mapped uniquely to the mitochondrial scaffold over at least 90% of their length were retained.

To obtain insight into the possible function or origins of the predicted ORFs, the putative amino acid sequence of each ORF was searched against the NCBI nr database with BLASTp using the BLOSUM45 matrix. Transmembrane domains were predicted with TMHMM v2.0 (Krogh et al. 2001) and secondary structural was predicted with SWISS-MODEL (Waterhouse et al. 2018).

All standard genes and predicted ORFs were indexed and quantified in terms of their relative expression dynamics with Kallisto (Bray et al. 2016) using each of the 18 developmental RNA-seq datasets used for gene annotation. Expression dynamics were visualised using the gplots (v3.0.1.1, Warnes et al. 2015), ggplot2 (v3.2.0, Wickham 2016) and scales (v1.0.0, Wickham and Seidel 2019) R libraries (v3.6.2, R Core Team 2020).

To identify genes or gene fragments that may have been transferred from the mitochondrial to the nuclear genome, often referred to as Nuclear Mitochondrial DNA (NUMT), a tBLASTn search was performed for each gene against the nuclear genome followed by manual inspection of the results. Positive hits were BLASTn searched against the NCBI nr database using the discontiguous megablast option to confirm mitochondrial affinity.

### Data availability

The data underlying this article are available from the NCBI Sequence Read Archive (SRA) database at https://www.ncbi.nlm.nih.gov/sra/and can be accessed with the BioProject ID PRJNA666063. The mitochondrial genome assembly is available with NCBI accession number MW080914.

## Acknowledgements

We thank the Vienna BioCenter Core Facilities (VBCF) next generation sequencing facility for their help with library construction and sequencing. This work was supported by the FWF Austrian Science Fund (grant number P 29455-B29 to AW).

## Authors’ contributions

ADC conceived of the study, designed the experiments, conducted the analyses and contributed to manuscript preparation. CB performed all laboratory work including DNA extractions. AW contributed to the design of the project and manuscript preparation.

## References

Abdelnoor RV, Christensen AC, Mohammed S, Munoz-Castillo B, Moriyama H, Mackenzie SA. 2006. Mitochondrial genome dynamics in plants and animals: convergent gene fusions of a MutS homologue. J. Mol. Evol. 63:165–173.

Alexeyev M, Shokolenko I, Wilson G, LeDoux S. 2013. The maintenance of mitochondrial DNA integrity - critical analysis and update. Cold Spring Harb. Perspect. Biol. 5:a012641.

Alves JMP, de Oliveira AL, Sandberg TOM, Moreno-Gallego JL, de Toledo MAF, de Moura EMM, Oliveira LS, Durham AM, Mehnert DU, Zanotto PM de A, et al. 2016. GenSeed-HMM: A Tool for Progressive Assembly Using Profile HMMs as Seeds and its Application in Alpavirinae Viral Discovery from Metagenomic Data. Front. Microbiol. 7:269.

Benson G. 1999. Tandem repeats finder: a program to analyze DNA sequences. Nucleic Acids Res. 27:573–580.

Boore JL, Medina M, Rosenberg LA. 2004. Complete sequences of the highly rearranged molluscan mitochondrial genomes of the Scaphopod Graptacme eborea and the bivalve Mytilus edulis. Mol. Biol. Evol. 21:1492–1503.

Bray NL, Pimentel H, Melsted P, Pachter L. 2016. Near-optimal probabilistic RNA-seq quantification. Nat. Biotechnol. 34:525–527.

Breton S, Beaupré HD, Stewart DT, Hoeh WR, Blier PU. 2007. The unusual system of doubly uniparental inheritance of mtDNA: isn’t one enough? Trends Genet. 23:465–474.

Breton S, Beaupré HD, Stewart DT, Piontkivska H, Karmakar M, Bogan AE, Blier PU, Hoeh WR. 2009. Comparative mitochondrial genomics of freshwater mussels (Bivalvia: Unionoida) with doubly uniparental inheritance of mtDNA: gender-specific open reading frames and putative origins of replication. Genetics 183:1575–1589.

Breton S, Milani L, Ghiselli F, Guerra D, Stewart DT, Passamonti M. 2014. A resourceful genome: updating the functional repertoire and evolutionary role of animal mitochondrial DNAs. Trends Genet. 30:555–564.

Calcino AD, de Oliveira AL, Simakov O, Schwaha T, Zieger E, Wollesen T, Wanninger A. 2019. The quagga mussel genome and the evolution of freshwater tolerance. DNA Res. 26:411–422.

Camacho C, Coulouris G, Avagyan V, Ma N, Papadopoulos J, Bealer K, Madden TL. 2009. BLAST+: architecture and applications. BMC Bioinformatics 10:421.

Chase CD. 2007. Cytoplasmic male sterility: a window to the world of plant mitochondrial-nuclear interactions. Trends Genet. 23:81–90.

Chen XJ. 2013. Mechanism of homologous recombination and implications for aging-related deletions in mitochondrial DNA. Microbiol. Mol. Biol. Rev. 77:476–496.

Christensen AC. 2013. Plant mitochondrial genome evolution can be explained by DNA repair mechanisms. Genome Biol. Evol. 5:1079–1086.

Christensen AC. 2014. Genes and Junk in Plant Mitochondria - Repair Mechanisms and Selection. Genome Biol. Evol. 6:1448–1453.

Combosch DJ, Collins TM, Glover EA, Graf DL, Harper EM, Healy JM, Kawauchi GY, Lemer S, McIntyre E, Strong EE, et al. 2017. A family-level Tree of Life for bivalves based on a Sanger-sequencing approach. Mol. Phylogenet. Evol. 107:191–208.

Cragg SA. 1996. The phylogenetic significance of some anatomical features of bivalve veliger larvae. In: Taylor JD, editor. Origin and Evolutionary Radiation of the Mollusca. Oxford University Press. p. 371–380.

Davila JI, Arrieta-Montiel MP, Wamboldt Y, Cao J, Hagmann J, Shedge V, Xu Y-Z, Weigel D, Mackenzie SA. 2011. Double-strand break repair processes drive evolution of the mitochondrial genome in Arabidopsis. BMC Biol. 9:64.

Donath A, Jühling F, Al-Arab M, Bernhart SH, Reinhardt F, Stadler PF, Middendorf M, Bernt M. 2019. Improved annotation of protein-coding genes boundaries in metazoan mitochondrial genomes. Nucleic Acids Res. 47:10543–10552.

Doucet-Beaupré H, Breton S, Chapman EG, Blier PU, Bogan AE, Stewart DT, Hoeh WR. 2010. Mitochondrial phylogenomics of the Bivalvia (Mollusca): searching for the origin and mitogenomic correlates of doubly uniparental inheritance of mtDNA. BMC Evol. Biol. 10:50.

Edwards JA, Edwards RA. 2019. Fastq-pair: efficient synchronization of paired-end fastq files. bioRxiv [Internet]:552885. Available from: https://www.biorxiv.org/content/10.1101/552885v1

Giribet G. 2008. Bivalvia. In: Ponder W, Lindberg DRR, editors. Phylogeny and Evolution of the Mollusca. University of California Press: Berkeley. p. 105–141.

González VL, Andrade SCS, Bieler R, Collins TM, Dunn CW, Mikkelsen PM, Taylor JD, Giribet G. 2015. A phylogenetic backbone for Bivalvia: an RNA-seq approach. Proc. Biol. Sci. 282:20142332.

Hanson MR, Bentolila S. 2004. Interactions of mitochondrial and nuclear genes that affect male gametophyte development. Plant Cell 16 Suppl:S154–S169.

Harzhauser M, Mandic O. 2010. Neogene dreissenids in Central Europe: evolutionary shifts and diversity changes. In: van der Velde Sanjeevi Rajagopal Abraham bij de Vaate G, editor. The Zebra Mussel in Europe. Weikersheim, Germany: Margraf Publishers GmbH. p. 11–29.

Heiler K, bij de Vaate A, Ekschmitt K, von Oheimb P, Albrecht C, Wilke T. 2013. Reconstruction of the early invasion history of the quagga mussel (Dreissena rostriformis bugensis) in Western Europe. AI 8:53–57.

Jackman SD, Coombe L, Warren RL, Kirk H, Trinh E, MacLeod T, Pleasance S, Pandoh P, Zhao Y, Coope RJ, et al. 2020. Complete mitochondrial genome of a gymnosperm, Sitka spruce (Picea sitchensis), indicates a complex physical structure. Genome Biol. Evol. 12:1174–1179.

Katoh K, Standley DM. 2013. MAFFT multiple sequence alignment software version 7: improvements in performance and usability. Mol. Biol. Evol. 30:772–780.

Kong L, Li Y, Kocot KM, Yang Y, Qi L, Li Q, Halanych KM. 2020. Mitogenomics reveals phylogenetic relationships of Arcoida (Mollusca, Bivalvia) and multiple independent expansions and contractions in mitochondrial genome size. Mol. Phylogenet. Evol. 150:106857.

Koren S, Walenz BP, Berlin K, Miller JR, Bergman NH, Phillippy AM. 2017. Canu: scalable and accurate long-read assembly via adaptive k-mer weighting and repeat separation. Genome Res. 27:722–736.

Kozik A, Rowan BA, Lavelle D, Berke L, Schranz ME, Michelmore RW, Christensen AC. 2019. The alternative reality of plant mitochondrial DNA: One ring does not rule them all. PLoS Genet. 15:e1008373.

Krogh A, Larsson B, von Heijne G, Sonnhammer EL. 2001. Predicting transmembrane protein topology with a hidden Markov model: application to complete genomes. J. Mol. Biol. 305:567–580.

Ladoukakis ED, Theologidis I, Rodakis GC, Zouros E. 2011. Homologous recombination between highly diverged mitochondrial sequences: examples from maternally and paternally transmitted genomes. Mol. Biol. Evol. 28:1847–1859.

Ladoukakis ED, Zouros E. 2001. Direct evidence for homologous recombination in mussel (Mytilus galloprovincialis) mitochondrial DNA. Mol. Biol. Evol. 18:1168–1175.

Li H. 2013. Aligning sequence reads, clone sequences and assembly contigs with BWA-MEM. 1303.3997 [Internet]. Available from: http://github.com/lh3/bwa

Li H. 2018. Minimap2: pairwise alignment for nucleotide sequences. Bioinformatics 34:3094–3100.

Liu Y-G, Kurokawa T, Sekino M, Tanabe T, Watanabe K. 2013. Complete mitochondrial DNA sequence of the ark shell Scapharca broughtonii: an ultra-large metazoan mitochondrial genome. Comp. Biochem. Physiol. Part D Genomics Proteomics 8:72–81.

Li W, Godzik A. 2006. Cd-hit: a fast program for clustering and comparing large sets of protein or nucleotide sequences. Bioinformatics 22:1658–1659.

McCartney MA, Auch B, Kono T, Mallez S, Zhang Y, Obille A, Becker A, Abrahante JE, Garbe J, Badalamenti JP, et al. 2019. The Genome of the Zebra Mussel, Dreissena polymorpha: A Resource for Invasive Species Research. bioRxiv [Internet]:696732. Available from: https://www.biorxiv.org/content/10.1101/696732v1

Mills EL. 1993. Colonization, Ecology, and Population Structure of the “Quagga” Mussel (Bivalvia: Dreissenidae) in the Lower Great Lakes. Can. S. Fish. Aquar. Sci., Vo 1:50.

Passamonti M, Ghiselli F. 2009. Doubly uniparental inheritance: two mitochondrial genomes, one precious model for organelle DNA inheritance and evolution. DNA Cell Biol. 28:79–89.

Plazzi F, Passamonti M. 2019. Footprints of unconventional mitochondrial inheritance in bivalve phylogeny: Signatures of positive selection on clades with doubly uniparental inheritance. J. Zoolog. Syst. Evol. Res. 57:258–271.

Quinlan AR, Hall IM. 2010. BEDTools: a flexible suite of utilities for comparing genomic features. Bioinformatics 26:841–842.

R Core Team. 2020. R: A Language and Environment for Statistical Computing. Available from: https://www.R-project.org/

Robertson G, Schein J, Chiu R, Corbett R, Field M, Jackman SD, Mungall K, Lee S, Okada HM, Qian JQ, et al. 2010. De novo assembly and analysis of RNA-seq data. Nat. Methods 7:909–912.

Smith DR. 2020. Can Green Algal Plastid Genome Size Be Explained by DNA Repair Mechanisms? Genome Biol. Evol. 12:3797–3802.

Soroka M, Rymaszewska A, Sanko T, Przylucka A, Lubosny M, Smietanka B, Burzynski A. 2018. Next-generation sequencing of Dreissena polymorpha transcriptome sheds light on its mitochondrial DNA. Hydrobiologia 810:255–263.

Stampar SN, Broe MB, Macrander J, Reitzel AM, Brugler MR, Daly M. 2019. Linear Mitochondrial Genome in Anthozoa (Cnidaria): A Case Study in Ceriantharia. Sci. Rep. 9:6094.

Warnes GR, Bolker B, Bonebakker L, Gentleman R, Liaw WHA, Lumley T, Maechler M, Magnusson A, Moeller S, Schwartz M, et al. 2015. gplots: Various R programming tools for plotting data. Available from: https://www.scienceopen.com/document?vid=0e5d8e31-1fe4-492f-a3d8-8cd71b2b8ad9

Waterhouse A, Bertoni M, Bienert S, Studer G, Tauriello G, Gumienny R, Heer FT, de Beer TAP, Rempfer C, Bordoli L, et al. 2018. SWISS-MODEL: homology modelling of protein structures and complexes. Nucleic Acids Res. 46:W296–W303.

Watters GT. 1999. Morphology of the Conglutinate of the Kidneyshell Freshwater Mussel, Ptychobranchus fasciolaris. Invertebr. Biol. 118:289–295.

Wickham H. 2016. ggplot2: Elegant Graphics for Data Analysis. Available from: https://ggplot2.tidyverse.org

Wickham H, Seidel D. 2019. scales: Scale Functions for Visualization. Available from: https://CRAN.R-project.org/package=scales

Wu X, Li X, Li L, Xu X, Xia J, Yu Z. 2012. New features of Asian Crassostrea oyster mitochondrial genomes: a novel alloacceptor tRNA gene recruitment and two novel ORFs. Gene 507:112–118.

Zouros E. 2013. Biparental Inheritance Through Uniparental Transmission: The Doubly Uniparental Inheritance (DUI) of Mitochondrial DNA. Evol. Biol. 40:1–31.

Zouros E, Ball AO, Saavedra C, Freeman KR. 1994. Mitochondrial DNA inheritance. Nature 368:818–818.

