## Supplementary Figures for "Heteroplasmy and repeat expansion in the plant-like mitochondrial genome of a bivalve mollusc"

### The expanded plant-like mitogenome of the invasive quagga mussel, *Dreissena rostriformis* - supplementary figures

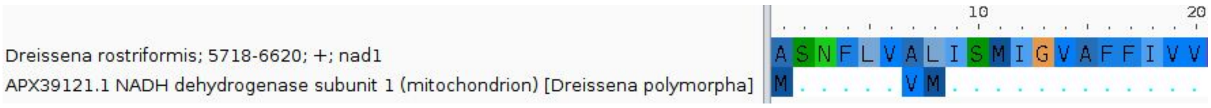


**Figure S1.** Alignment of the first 20 amino acids of the zebra mussel NAD1 (bottom, APX39121.1) with the corresponding region from the quagga mussel (top) with only differences displayed in the zebra mussel track. As no methionine was identified in the quagga mussel that corresponded to the two at positions 1 and 8 in the zebra mussel, the position 11 methionine was selected as the likely start site of the quagga mussel NAD1.

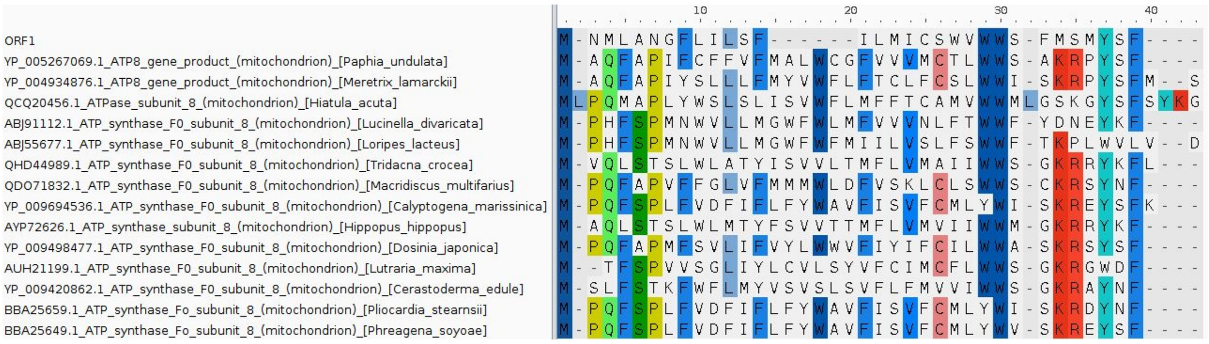


**Figure S2.** Alignment of the putative atp8 from the quagga mussel (ORF1) with other Imparidentia atp8 orthologues. Majority rule consensus sites are highlighted.

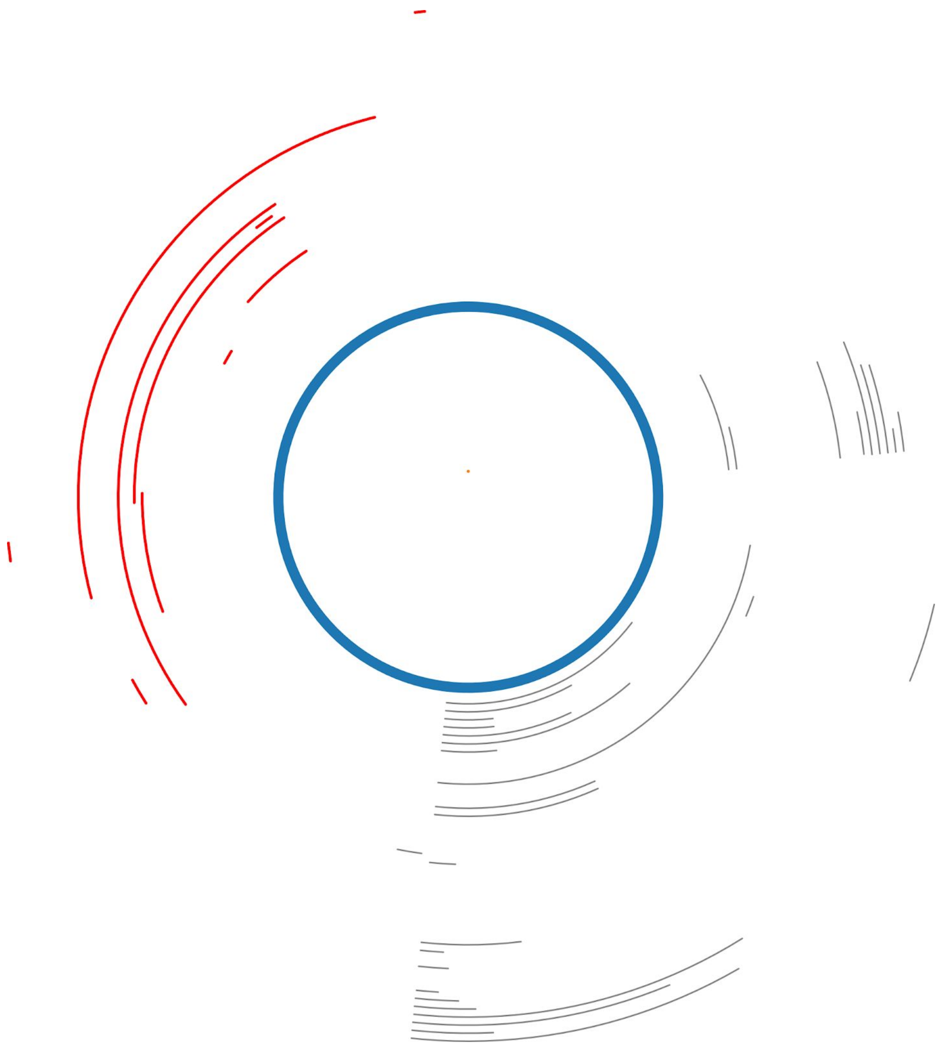

**Figure S3.** Clipped ends mapped back to the genome assembly and visualised with ConcatMap. The blue circle represents the genome, the grey lines represent the clipped ends that mapped back to coding regions and the red lines represent the clipped ends that mapped back to the large repetitive region. It was full length reads corresponding to the red mapped clipped ends that were further investigated for repeat content.

a)

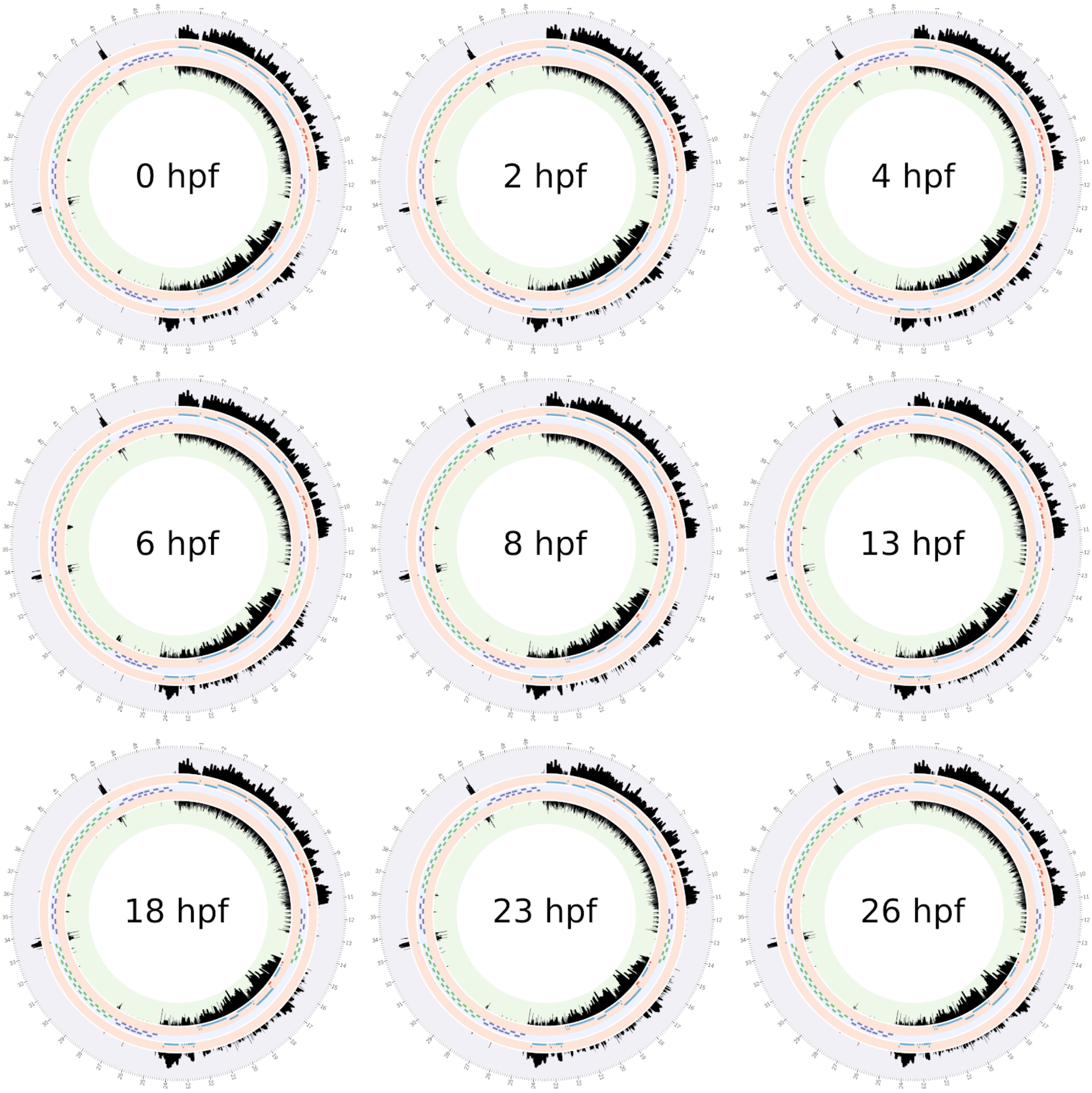

b)

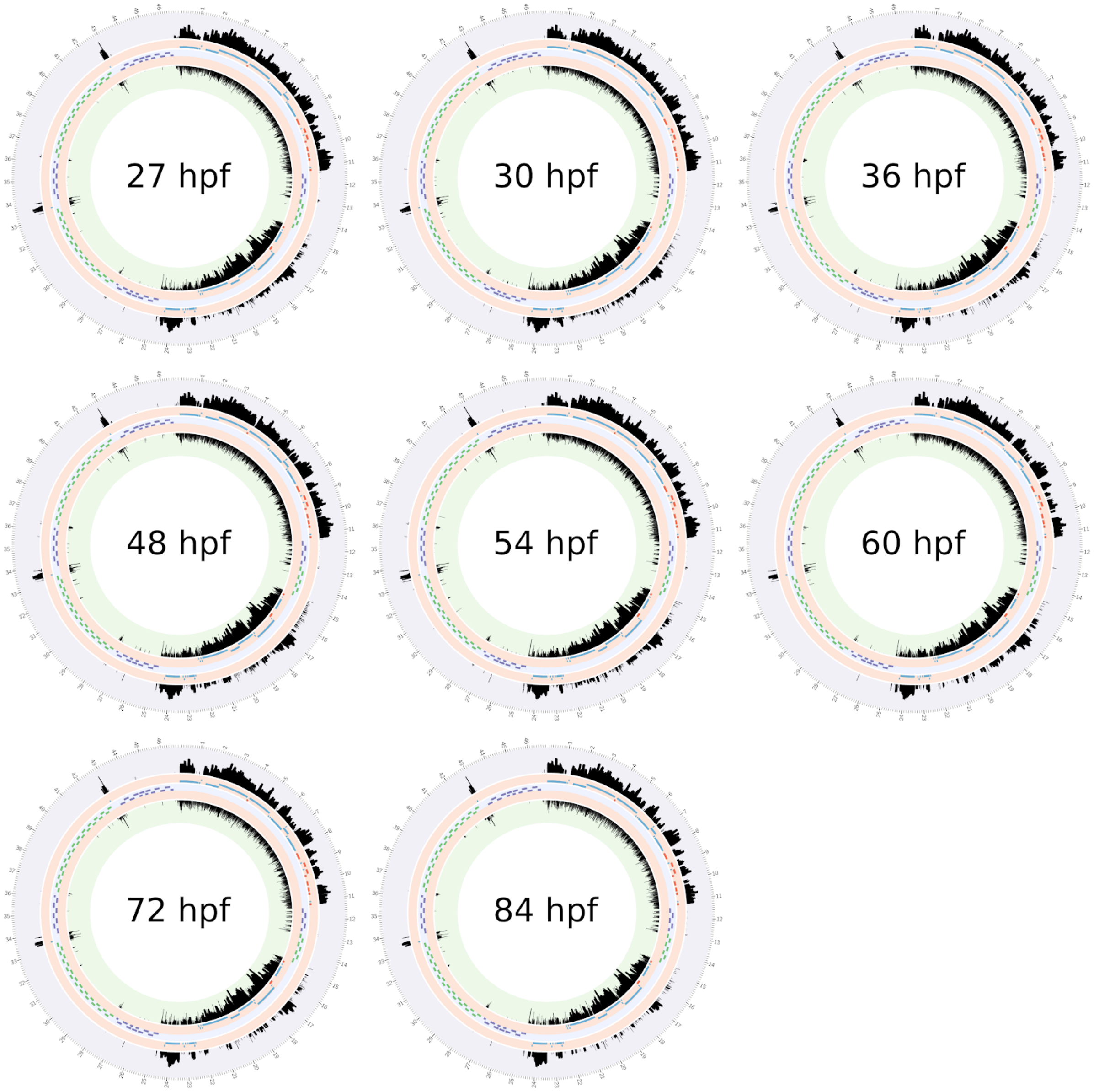


**Figure S4.** RNA-seq coverage. Each circos plot represents a developmental stage with the inner and outer tracks displaying the log of the reads per million of mapped reads (RPM) RNA-seq coverage values values for the light and heavy strands respectively. The middle track shows the locations of the 308 bp (purple) and 258 bp (green) tandem repeats with the remaining two tracks showing the location of coding and ribosomal genes (long, blue), tRNAs (short, blue) and novel ORFs (red) mapping to the light and heavy strand, respectively. Reads were allowed to map to multiple locations to avoid the loss of reads that may have been derived from repetitive loci. Stages include unfertilised eggs (0 hpf), 2-4 cell embryos (2 hpf), gastrula (4 hpf), swimming gastrula (6 hpf - 8 hpf), trochophores (13 hpf - 23 hpf), early veligers (26 hpf - 30 hpf), D-shaped veligers (36 hpf - 72 hpf) and late-stage veligers (84 hpf).
