## Supplementary Methods for "Heteroplasmy and repeat expansion in the plant-like mitochondrial genome of a bivalve mollusc"

**The expanded plant-like mitogenome of the invasive quagga mussel, *Dreissena rostriformis* - supplementary methods**

**Genome assembly**

```
#correct pacbio reads with canu
canu -s spec.conf -p Dro -d /path/to/output -pacbio-raw /path/to/raw/pacbio/reads/reads.fa

#canu spec.conf
genomeSize=1.6g
batMemory=100g
batThreads=20
correctedErrorRate=0.15
corOutCoverage=200

#Run genseed
genseed-hmm_26b.pl -conf config.txt

#genseed config file
seed = seed_COI.fa
db = canu_corrected_pacbio_reads.fa
assembler_name = cap3
next_seed_size = 40
blastn_parameters = -evalue 1e-15 -num_threads 1 -perc_identity 98
-max_target_seqs 50000 -dust no
output = genseed_canu_corrected
clean = no

#Map uncorrected pacbio reads back to genseed assembly
pbmm2 align --best-n 1 --preset "SUBREAD" --sort --log-level "DEBUG"
--min-concordance-perc 0 Dro_genseed_mito.mmi pacbio_reads.fofn
Dro_genseed_mito.aligned.bam -j 16

#Run canu with these reads
canu -s spec.conf -p Dro_mito -d /path/to/output/canu_genseed
/path/to/mito/reads/Dro_genseed_mito.aligned.fastq.gz

#canu spec.conf
genomeSize=100k
batMemory=20g
batThreads=16
correctedErrorRate=0.040
corOutCoverage=1000
corMaxEvidenceErate=0.15

#Polish assembly with gcpp using all subreads
```

```
pbmm2 align --preset "SUBREAD" -j 8 --sort --log-level "DEBUG"
Dro_mito.contig.mmi pacbio_reads.fofn Dro_mito.contig.bam
gcpp -r Dro_mito.contig.fa -o
Dro.mito.corrected.fasta,Dro.mito.corrected.fastq,Dro.mito.corrected.gff
--annotate-gff -j 8 Dro_mito.contig.bam
```

```
#Run ConcatMap to visualise mapping of all subreads to assembly after trimming
down canu assemly so it only includes one copy of the genome and making sure that
the genome assembly is repeated so that there are two copies in tandem.

ConcatMap.py  -i aln.sam -r 47454 -f pdf -w 0.01 -c 10 -s 50
```

**Transcriptomics**

```
#Build de-novo transcriptomes for all developmental datasets
transabyss --pe lib1_1.fq lib1_2.fq --SS --length 150 --threads 8 --outdir ./

#Combine all de-novo transcriptomes and deduplicate
for i in `cat names.txt` ; do cat "$i"/transabyss-final_rev.fa | sed -e "s/>/>$i\_/" -e 's/\
/\_/g' >>total_transabyss-final_rev.fa ; done
cd-hit -T 12 -M 2000 -i total_transabyss-final_rev.fa -o
total_transabyss-final_deduped.fa -c 0.98

#Map total_transabyss-final_deduped.fa to unmasked mitochondria-free genome
minimap2 -d Dro_genome.mmi Dro_genome.fa
minimap2 -ax splice -C5 --secondary=no -uf -t 12 --cs Dro_genome.mmi
total_transabyss-final_deduped.fa >total_transabyss-final_deduped_aligned.sam
minimap2 -cx splice -C5 --secondary=no -uf -t 12 --cs Dro_genome.mmi
total_transabyss-final_deduped.fa >total_transabyss-final_dedulped_aligned.paf

#Pull mito mapping entries that map at least 90% of their length and check that they
only map to a single location in the genome
awk '{ if (($4-$3) >= (0.9*$2)) {print $0} }' total_transabyss-final_deduped_aligned.paf
| grep tig00000001 >mito_mapping_90+.paf
awk '{ if (($4-$3) >= (0.9*$2)) {print $0} }' total_transabyss-final_deduped_aligned.paf
| cut -f1 | sort | uniq -c | grep '\ 1\ ' | sed 's/\ \ \ \ \ \ 1\ //' >uniqes.names
fgrep -f uniqes.names mito_mapping_90+.paf >mito_mapping_90+_uniques.paf

#Convert paf to bed
paftools.js splice2bed mito_mapping_90+_uniques.paf
>mito_mapping_90+_uniques.bed

#Convert bed to gff3 with genometools
gt bed_to_gff3 -o mito_mapping_90+_uniques.gff mito_mapping_90+_uniques.bed
```

#To get fasta of mapped reads from bed file
fastaFromBed -fi Dro_genome.fa -name -s -split -bed mito_mapping_90+_uniques.bed >mito_mapping_90+_uniques.fa

#Convert gff to gtf that Transdecoder likes
grep -P -v '#' mito_mapping_90+_uniques.gff | sed 's/BED_feature/transcript/g' | sed 's/BED_block/exon/' | grep -v BED_thick_feature | sed 's/ID=/gene_id\ \"/' | sed 's/;Name=/\";\ transcript_id\ \"/' | sed 's/$/\";/' | sed 's/Parent=/gene_id\ "/' >mito_mapping_90+_uniques.gtf

#Produce a fasta file from this gtf file with the appropriate headers
gtf_genome_to_cdna_fasta.pl mito_mapping_90+_uniques.gtf Dro_genome.fa >mito_mapping_90+_uniques_transdecoder.fa

#Re-convert the gtf into a gff3 file format that Transdecoder likes
gtf_to_alignment_gff3.pl mito_mapping_90+_uniques.gtf >mito_mapping_90+_uniques_transdecoder.gff3

#Deduplicate fasta file with cd-hit
cd-hit -T 12 -i mito_mapping_90+_uniques.fa -o mito_mapping_90+_uniques_deduped.fa -c 0.95

**GC content**

#Get GC content of sliding window
perl GC_content.pl --fasta Dro_genome.fa --window 40 --step 40

**Genome coverage**

#Map a short read library against the nuclear genome without the mitochondrial genome.
bwa index Dro_nuclear_genome.fa
bwa mem -t 16  Dro_nuclear_genome.fa illumina_1.fq illumina_2.fq | samtools view -bS - >aligned.bam

#Extract unmapped reads
samtools view -Sbh -f 4 aligned.bam >unaligned.bam
samtools view -Sh unaligned.bam >unaligned.sam

#Manually convert to fastq and sort output
cat unaligned.sam | grep -v ^@ | awk 'NR%2==1 {print "@"$1"\n"$10"\n+\n"$11}' | paste - - - - | sort -k1,1 -S 3G | tr '\t' '\n' >unaligned_1.fastq
cat unaligned.sam | grep -v ^@ | awk 'NR%2==0 {print "@"$1"\n"$10"\n+\n"$11}' | paste - - - - | sort -k1,1 -S 3G | tr '\t' '\n' > unaligned_2.fastq

#Extract just the paired reads
fastq_pair unaligned_1.fastq unaligned_2.fastq

#Map reads to mito genome
bwa mem -t 8 Dro_genome.fa unaligned_1.fastq.paired.fq
unaligned_2.fastq.paired.fq >Dro_genome_mapped.sam

#Convert sam to six field bed
sam2bed <Dro_genome_mapped.sam >Dro_genome_mapped.bed
cut -f1-6 Dro_genome_mapped.bed >Dro_genome_mapped_6field.bed

#Get bga coverage files
bedtools genomecov -bga -strand + -i Dro_genome_mapped_6field.bed -g
genome.txt >plus_strand_cov.bga
bedtools genomecov -bga -strand - -i Dro_genome_mapped_6field.bed -g
genome.txt >minus_strand_cov.bga

**Concatmap**

#Map long reads to genome (2x construct) with minimap2
minimap2 -t 12 -ax -N 0 map-pb Dro.genome.double.fa
/path/to/mito/reads/Dro_genseed_mito.aligned.fastq.gz
>aln_noSecondary.sam

#Run ConcatMap.py
python3 ./ConcatMap_v1.2.py -i aln_noSecondary.sam -o no_clip -r 46868 -m 5000
-f png -c 0.5 -l 0.02 -w 0.25 -s 20 -x t

#Convert sam to bam
samtools view -S -b aln_noSecondary.sam >aln_noSecondary.bam

#Convert bam to bed
bamtools convert -format bed -in aln_noSecondary.bam -out aln_noSecondary.bed

#Run with reads >40kb
python3 ./ConcatMap_v1.2.py -i aln_noSecondary.sam -r 46868 -m 40000 -f png -c
0.5 -l 0.02 -w 1 -s 20 -x t
